# MetaGLIMPSE: Meta Imputation of Low Coverage Sequencing Data for Modern and Ancient Genomes

**DOI:** 10.1101/2025.06.24.660721

**Authors:** Kiran H. Kumar, Simone Rubinacci, Sebastian Zöllner

## Abstract

The advent of efficient and accurate imputation for low coverage sequencing offers an unbiased alternative to SNP array imputation, increasing the accuracy of rare variant imputation across all populations. Since imputation accuracy generally increases with larger reference panels and closer ancestry match between target and reference samples, leveraging imputation from multiple reference panels improves imputation accuracy; however, individual reference panel genotypes are often privacy protected. Meta-imputation bypasses individual level data by combining single-panel imputed genotypes through estimating panel and marker specific weights. We present a novel meta-imputation method, MetaGLIMPSE, that combines estimates from multiple reference panels for low coverage sequencing imputation. Across all our scenarios, for both modern and ancient DNA samples, MetaGLIMPSE consistently outperforms the best single panel imputation for coverages 0.1x - 8x and across all minor allele frequencies, equaling the combined panel imputation for some parameters. Finally, MetaGLIMPSE is computationally efficient, meta-imputing 500 whole genomes in 16% of the time of GLIMPSE2.

## Introduction

Genomic imputation is a cost-effective tool that powers downstream analysis such as GWAS, fine mapping, and meta-analyses (Browning and Browning 2007; Das et al. 2018; Howie et al. 2012; Fuchsberger et al. 2015; Li et al. 2010; Marcini et al. 2010; Yu et al. 2022). Earlier algorithms focused on SNP array data (Browning and Browning et al. 2007; Fuchsberger et al. 2015; Howie et al. 2012). Nowadays, low coverage sequencing is a growing alternative to SNP arrays due to decreasing sequencing costs (Boltz et al. 2024; Martin et al. 2021). Low coverage sequencing involves shotgun sequencing at low mean depths between 0.1x and 8x (Boltz et al. 2024; Martin et al. 2021; Rubinacci et al. 2021). While such read data is often insufficient to confidently call genotypes, imputation accuracy for low pass sequencing at coverages as low as 0.5x is comparable to commonly used SNP arrays, and a coverage of 1x suffices for confident rare variant detection. (Martin et al. 2021; Rubinacci et al. 2021). Moreover, low coverage sequencing is especially useful for non-European populations as it provides unbiased coverage across the genome and eliminates ascertainment bias inherent in the design of SNP arrays (Boltz et al. 2024; Martin et al. 2021), which facilitates downstream analyses such as calculating polygenic risk scores (Nguyen et al. 2025). This advantage has motivated large low coverage sequencing efforts, such as Blended Genome Exome (BGE), which has sequenced at least 53,000 samples at coverages between 1x to 4x (Boltz et al. 2024). Furthermore, the lower cost and experimental considerations make low-pass sequencing attractive for the sequencing of ancient DNA (aDNA); at least 30% of all aDNA samples are sequenced via low coverage. This would imply, very conservatively, that ∼3,000 aDNA low coverage genomes have been sequenced. This number is likely to be larger and grow exponentially in the coming decades (Mallick et al. 2024; Sousa da Mota et al. 2023).

The challenge of low coverage imputation is that individual genotypes are more uncertain, which precludes pre-phasing, a computation time saving strategy used in the SNP array imputation (Howie et al. 2012). Thus, an algorithm for low coverage imputation must handle both phasing and imputation with computational efficiency. A commonly used method for low coverage imputation for both modern and ancient DNA (aDNA) samples (Allentoft et al. 2024; Boltz et al. 2024; Erven et al. 2024; Nakamura et al. 2024; Ringbauer et al. 2024; Santos et al. 2024; Sousa da Mota et al. 2023) is GLIMPSE (Rubinacci et al. 2021) and its companion GLIMPSE2 (Rubinacci et al. 2022). GLIMPSE and GLIMPSE2 iteratively phase and impute using a Gibbs Sampler, resulting in phased genotypes.

Aside from algorithmic considerations, the main two determinants of imputation accuracy are ancestry match with target samples and reference panel size. The latter has a large benefit, in particular, on the imputation accuracy of rare variants, which are associated with substantial functional implications (Das et al. 2018). For this reason, the size of reference panels has grown over time, the largest ones broadly used are TopMed and Human Reference Consortium (HRC) representing 133,597 (Smith et al. 2023), and 32,470 (Das et al. 2016; McCarthy et al. 2016; Taliun et al. 2021) individuals. However, for underrepresented populations imputing with an ancestry specific panel may result in more accurate genotypes than imputing with a large, primarily European ancestry reference panel (Carlson et al. 2025; Das et al. 2018; Jewett et al. 2012; Yu et al. 2022). The optimal method leverages genotypes from multiple reference panels by combining the smaller panels into a ‘mega’ panel and then imputing once with this mega panel. However, creating such a panel is often not possible because the individual level data is privacy protected. Furthermore, merging two reference panels, which are genotyped and quality controlled using different algorithms and parameters, is not trivial, as seen in the creation of the HGDP+1KG panel (Koenig et al. 2024). Finally, combining a small population-specific reference panel with a large nonspecific panel may cause rare population-specific haplotypes to be ignored in favor of more frequent haplotypes, reducing imputation accuracy. To address this challenge for imputation from individual-level genotyping data, Yu et al. 2022 proposed a “meta-imputation” method, MetaMinimac2, that combines imputation results generated from chip-genotypes with multiple reference panels. Their method has been applied to improve imputation accuracy for multiple ancestries, including South Asian samples (Li et al. 2025). However, MetaMinimac2 cannot be applied to low coverage sequencing data, as the input data is assumed to be phased genotypes without uncertainty.

In this paper, we introduce the first method for meta-imputation for low coverage sequencing data, called MetaGLIMPSE, which enables imputation of low coverage sequencing data with multiple reference panels. Through meta imputation, we bypass accessing individual reference panel samples. We construct a Hidden Markov Model (HMM) to estimate weights, used to directly combine single-panel imputation results into a consensus genotype.

We evaluate our method on high coverage sequencing data from 1000 Genomes for 4 scenarios, down sampling to 0.1x to 8x. First, we evaluate the case in which the target samples and reference panels originate from the same ancestry, modeling combining imputation results from two reference panels that have the same ancestry as the target sample. Second, we evaluate the case in which the target samples are an admixture of the ancestries in each reference panel, modeling imputing admixed individuals with two reference panels, each containing a different source population. Third, we evaluate the case where the target samples are aDNA and the reference panels are from two different modern populations. In all these cases, our method outperforms the best-available single reference panel imputation across all minor allele frequencies (MAFs). In summary, MetaGLIMPSE delivers imputed genotypes that are more accurate than single panel imputation at standard mean coverages (0.1x - 8x) without knowing, *a priori*, which reference panel is the best.

## Subjects and Methods

MetaGLIMPSE combines multiple low-coverage sequencing imputation results by calculating each imputation result’s fit with the sequencing data. The imputation results, along with the genotype likelihoods, are inputs to a HMM that estimates weights for each panel at every imputed polymorphism. We leverage these weights to combine the single-panel imputation estimates into meta-imputed genotypes.

## Model Description

In a dataset where variants have been called at *M* loci, we model sequencing data for each individual as genotype likelihoods *GL*_*m*_ = *[GL(0), GL(1), GL(2)]* at each called variant *m*. Let there be *K* distinct reference panels, and let *(A*_*1,k,m*_, *A*_*2,k,m*_) denote the imputation output for marker *m* (1 ≤ *m* ≤ *M*) with panel *k* (1 ≤ *k* ≤ *K*). *(A*_*1,k,m*_, *A*_*2,k,m*_) are the outputted phased dosages of any low coverage sequencing algorithm, e.g. GLIMPSE2. Therefore, *A*_*1,k,m*_ + *A*_*2,m*_ is the unphased imputed genotype dosage for marker *m* and panel *k*. Finally, let *G*_*m*_: 0 ≤ *G*_*m*_ ≤ 2 be the meta-imputed genotypes, obtained through estimating marker and panel-pair specific weights via HMM as described in the following sections.

## Defining the HMM

### Hidden and Observed States

We define an HMM across all imputed genotypes *m* where the observed states are the genotype likelihoods *GL*_*m*_, and the hidden states are the “best” imputation results, denoted *A*^*\**^, = *(A* ^*\*,p1, m*^, *A* _^*\**^, *p2, m*_): A^*^[i] ∈{*A*_*k, pj, m*_: 1 ≤ *k* ≤ *K*; *p*_*j*_ ∈{1,2}; 1 ≤ *m* ≤ *M*}, where *p*_*j*_ represents phasing status.

### Emission and Transition Probabilities

The emission probability captures how well each imputation result fits with the observed sequencing data (Equation 1).

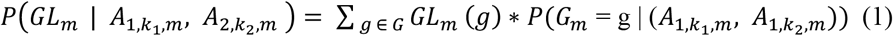

Due to the genotype uncertainty at each marker, we marginalize over the possible unphased genotypes in the emission probability. We also sum over phase since the imputation results are phased, whereas the genotype likelihoods are not. The transition probability is a function of marker distance based on the Li and Stephens transition probabilities for diploid data (Ausmees et al. 2023; Li et al. 2010).

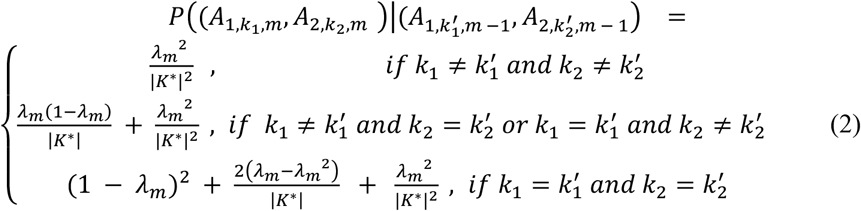

Here we assume that the switch from one imputation result being optimal to a different imputation result being optimal (i.e. (*A*_1,∗ *m*-1_, *A*_2,∗,*m-*1_) ≠ (*A*_1,∗,*m*_, *A*_2,∗,*m*_)) is driven by the recombination process. Thus, λ_*m*_ = 1 - e^-*θ* * d(*m-1*, *m*)^, and *θ* = 2 ^*^ 10^-7^ is the recombination rate between the variants (Yu et al. 2022). *d(m-1, m)* is the base pair distance between the *m*^th^ and *m-1*^*th*^ markers. The probability of a transition between distinct hidden states increases as the distance between adjacent markers increases. We fix *θ* according to values found in the meta-imputation literature (Yu et al. 2022).

### Weight Estimation and Meta Imputed Genotypes

We apply the forward-backward algorithm (Baum 1970) to this HMM, generating the marginal probability of each imputation panel pair being optimal at variant *m*. These probabilities serve as the weights, denoted as 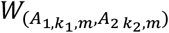 (Equation 3), where the subscripts indicate the reference panels *k*_*1*_ and *k*_*2*_ used for the two haplotypes.

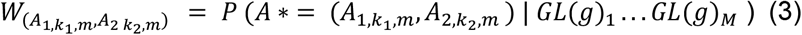

Using these weights, we compute the meta-imputed genotype at each marker by averaging the imputed genotypes from all possible panel pairs, weighted by their probabilities. Formally, for each marker *m*, we calculate *G*_*m*_ as follows: (Equation 4).

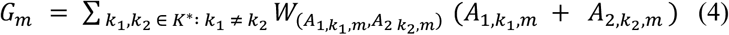

This derives an expected genotype value that reflects a consensus across all panels, giving more weight to the panels that best fit the observed sequencing data.

### Leveraging phasing information

MetaGLIMPSE can leverage the phase information on imputed genotypes by modifying the space of hidden states in the HMM. We describe two versions of MetaGLIMPSE. The default version, MetaGLIMPSE, leverages phasing, whereas “MetaGLIMPSE-plain” does not.

Formally, let 𝒮 be the set of all hidden states. For MetaGLIMPSE-plain, 𝒮={*(A*_*11*_,*A*_*12*_*),(A*_*21*_,*A*_*22*_),…,*(A*_*k1*_,*A*_*k2*_)}, requiring that *A*_^*\**^*1*_ and *A*_^*\**^*2*_ originate from the same reference panel. For the default MetaGLIMPSE, we define 𝒮={(*A*_*1i*_,*A*_*2j*_) : 1 ≤ *i,j* ≤ *k*}, thus allowing *A*_^*\**^*1*_ to originate from a different reference panel than A_^*^2_. MetaGLIMPSE-plain has |𝒮| = *K* hidden states, whereas MetaGLIMPSE has 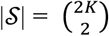 hidden states.

### Evaluation

To assess the performance of MetaGLIMPSE and MetaGLIMPSE-plain in a variety of scenarios we impute variants on chromosome 20 from the unrelated 2504 samples in 1000 Genomes Phase 3 resequenced at 30x (Byrska-Bishop et al. 2022) as the source dataset for both modern target samples and reference panels. We also evaluate ten target aDNA samples, a subset of samples described in Sousa da Mota et al. (2023) (Supplementary Table 1). We chose these ten for spatial diversity, with five Europeans, three Africans, and two Native-Americans and temporal diversity, as the samples range from 327 to 35,154 years before present. These samples also have relatively low C to T deamination rates. The maximum percentage of C→ T transitions at reads’ ends across all samples is 18.5% and the mean is 4.7% and the minimum is 0.8%. Three of the ten samples do not have reported deamination rates (Sousa da Mota et al. 2023).

**Table 1:** Grid of computation times by number of samples (N) and number of markers (M) while running all the chunks in parallel.

|  | M = 900,000 | M = 1 Genome |
| --- | --- | --- |
| N= 1 | 4.42 seconds | 3 minutes 41 seconds |
| N= 50 | 2 minutes 50 seconds | 2 hours 21 minutes 51 seconds |
| N = 500 | 32 minutes 7 seconds | 26 hours 45 min 42 seconds |

To create our test datasets, we first align the aDNA fasta files to hg38, unless the hg38 aligned bams already exist. We use bwa-mem2 with the default settings, as recommended for aDNA by Xu et al. (2021). All 1000 Genomes samples include hg38 aligned bams. Then, from the 2504 total modern samples, we remove samples that have any parents in the dataset, using the pedigree file from Byrska Bishop et al. (2022), leaving 2495 total unrelated samples. From these 2495 remaining modern samples as well as the ten ancient samples, we generate low coverage target samples by downsampling to a range of mean coverages {0.1x, 0.5x, 1x, 2x, 4x, 6x, 8x} using samtools view v1.19.2. For each sample and coverage, we calculate genotype likelihoods using bcftools mpileup v1.19 at all biallelic sites for the entire 3202 individuals in 1000 Genomes (Byrska-Bishop et al. 2022). We call the genotypes of aDNA in the same manner as the modern samples since the choice of caller does not substantively impact imputation results (Sousa da Mota et al. 2023).

We create five reference panels by subsetting the samples for the following populations based on the 1000 Genomes ancestry classification: Europeans half A (n = 249), Europeans half B (n = 249), Europeans (n = 498), Africans (n = 599, excluding African Americans), and Africans + Europeans (n = 1098, excluding African Americans). We generate European half A and European half B by randomly sampling half of the full European panel. We remove all variants that are singletons across the 2495 samples. Within every reference panel, we remove triallelic SNPs, and then we remove SNPs with an allele count of 0. When imputing a target sample, we exclude that sample from the reference panel.

We impute the target African-American, and aDNA samples with downsampled coverage using the African, European, and African + European-mega panels. We impute the downsampled Europeans with each sample’s distinct European A, European B, and full European panels. In all three experiments, we impute in 6cM length chunks using GLIMPSE 2.0.1 software -AP option branch for 19 burn-in iterations and 1 main iteration for each reference panel described above. We then meta-impute the modern African-Americans, modern Europeans and the aDNA samples using their respective single panel imputations via MetaGLIMPSE.

To compare the performance of both single panel imputation, mega-imputation, and meta-imputation, we identify markers that are polymorphic in both single-panel imputations and, for these markers. This results in ∼344k and ∼281k variants that are polymorphic in both the Europeans half A and Europeans half B panel and both the African and European panel, respectively. We calculate the accordance of each imputation result with the high coverage genotypes. For the modern samples, we use the 30x genotypes called in 1000 Genomes. For the ancient samples, we call the validation genotypes using the “five filter approach” (Moreno-Mayar et al. 2018; Sousa da Mota et al. 2023).

To calculate imputation accuracy, we bin variants by MAF and determine the aggregated Pearson R^2^ between the high coverage genotypes and imputed genotype estimates in each bin. Aggregated R^2^ combines the true genotypes into one vector and the estimated genotypes into a second vector and calculates the correlation between these two vectors (Rubinacci et al. 2021). This metric accounts for the relatively small number of target samples we impute, since the Pearson R^2^ per marker is not defined when the variance of either the true or imputed genotypes is zero (Li et al. 2023). We use aggRSquare v1.0.0 to compute the aggregated R^2^, and we set the allele frequency of a variant to its value across all unrelated samples in the 30x resequenced Phase 3 1000 Genomes dataset.

## Results

We evaluate MetaGLIMPSE across sequencing coverages ranging from 0.1x to 8x, considering a variety of realistic scenarios with two available reference panels. These scenarios include instances where (i) target samples share ancestry with both reference panels; (ii) where admixed individuals are present, with each panel representing one ancestral background, and; (iii) where aDNA samples are imputed with modern reference panels.

For each scenario, we generate four distinct imputed datasets for the target samples: Initially, two imputed cohorts are generated, each using one of the available reference panels. A third cohort is then created through mega-imputation, where both panels are merged into a unified reference.

The fourth cohort is produced by applying MetaGLIMPSE to combine the two individually imputed cohorts.

### Meta Imputing with reference panels from the same population

A common use of meta-imputation is to combine imputation results from several reference panels, such as TopMed and HRC, that cannot be merged in a mega-imputation framework, due to privacy reasons. We model this challenge by splitting the unrelated 30x European samples (n = 499) in 1000 Genomes (Byrska-Bishop et al. 2022) into two reference panels, while excluding the target sample from either panel. We imputed each of our 499 target samples with its distinct set of three reference panels separately: European mega panel (n = 498), European half A (n = 249), and European half B (n = 249). We then combined the results from European half A imputation and European half B imputation using MetaGLIMPSE. We compared MetaGLIMPSE to the two half-panel imputations and the one mega-imputation.

Across all coverages, mega-imputation, unsurprisingly, achieves the highest overall aggregated R^2^, with high accuracy for common variants that rapidly declines as alleles become rarer. European panels half A and half B have an almost identical aggregated R^2^ that is consistently lower than the aggregated R^2^ of the mega-imputation, with larger differences for uncommon and rare variants. MetaGLIMPSE provides more accurate imputation results than either of the individual European panels across all MAFs and all sequencing depths (Supplementary Figures 3-4).

At 2x coverage, aggregated R^2^ values for all imputation scenarios are within 0.01 of each other for variants with MAF >1%. The difference in imputation quality increases between MAF 0.1% and 1%. For MAFs between 0.1% and 0.2%, the mega imputation maintains an R^2^ of 0.68, while the individual panels drop to 0.62. MetaGLIMPSE gives an R^2^ of 0.65, recovering 45% of the performance increase of mega-imputation compared to the best single panel imputation (Figure 2a-b). Phasing also plays an important role in MetaGLIMPSE. At 2x, for MAFs between 0.1% and 0.2%, MetaGLIMPSE performance drops to 0.64, a difference of 0.01, when we do not leverage phasing (Supplementary Figures 3-4).

**Figure 1.**
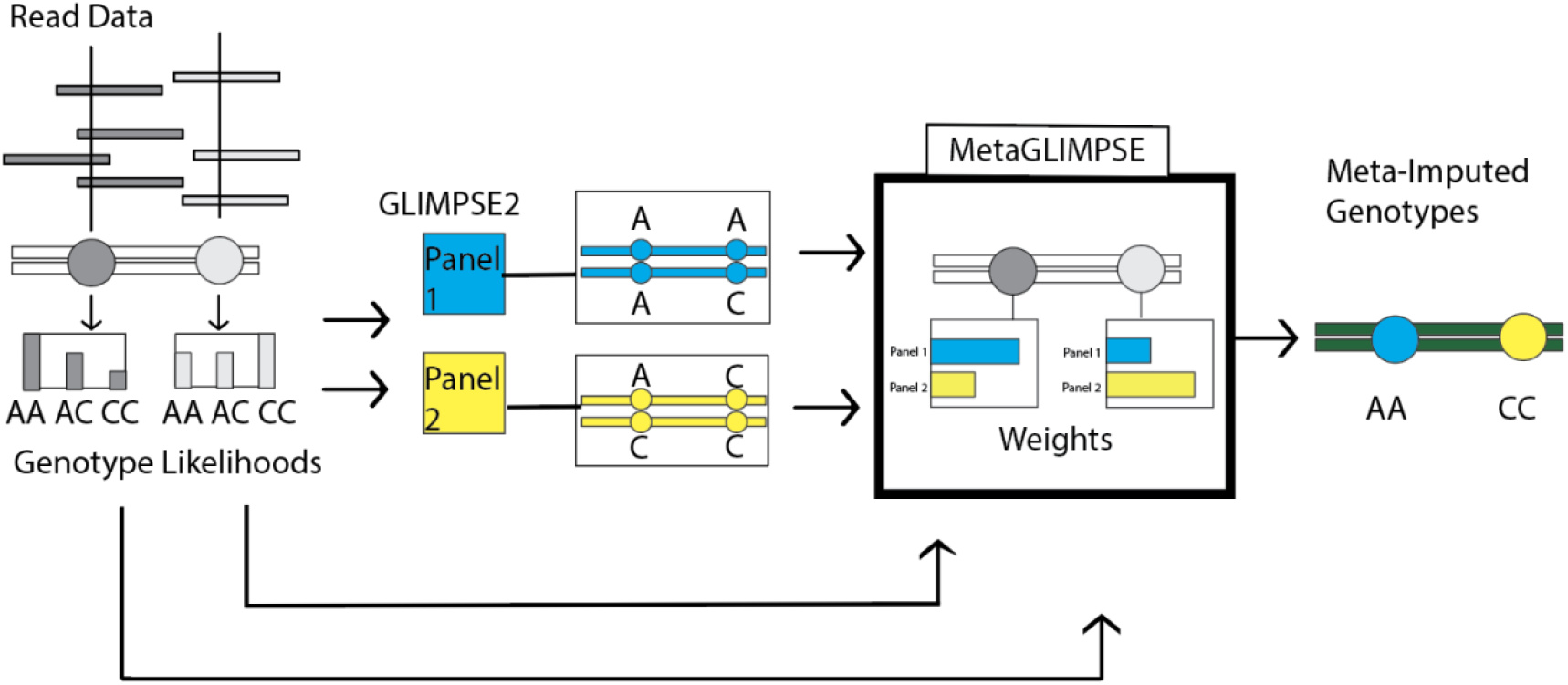
Model Overview of MetaGLIMPSE. The workflow for MetaGLIMPSE is shown here for a single target sample with two markers in dark and light grey, respectively. The workflow starts from the read data, from which genotypes likelihoods are called. Here, the most likely genotypes are A/A for the first marker and C/C for the second marker. Then, we phase and impute haplotypes based on the genotype likelihoods with two different reference panels separately: Panel 1 in blue, and Panel 2, in yellow via GLIMPSE2. The first panel imputes A|A, A|C, whereas the second panel imputes A|C, C|C. MetaGLIMPSE analyzes the imputed haplotypes and genotype likelihoods to estimate weights for each individual and marker, modelling the correlation of nearby markers by using a HMM. In this example, the first marker has a higher weight for panel 1 and the second marker has a higher weight for panel 2. Finally, MetaGLIMPSE outputs meta-imputed genotypes by generating a weighted average of each of the imputation estimates. In the example, MetaGLIMPSE imputes A\A, C\C as the best guess genotypes, which derive from the higher weights on panel 1 and panel 2, respectively.

**Figure 2.**
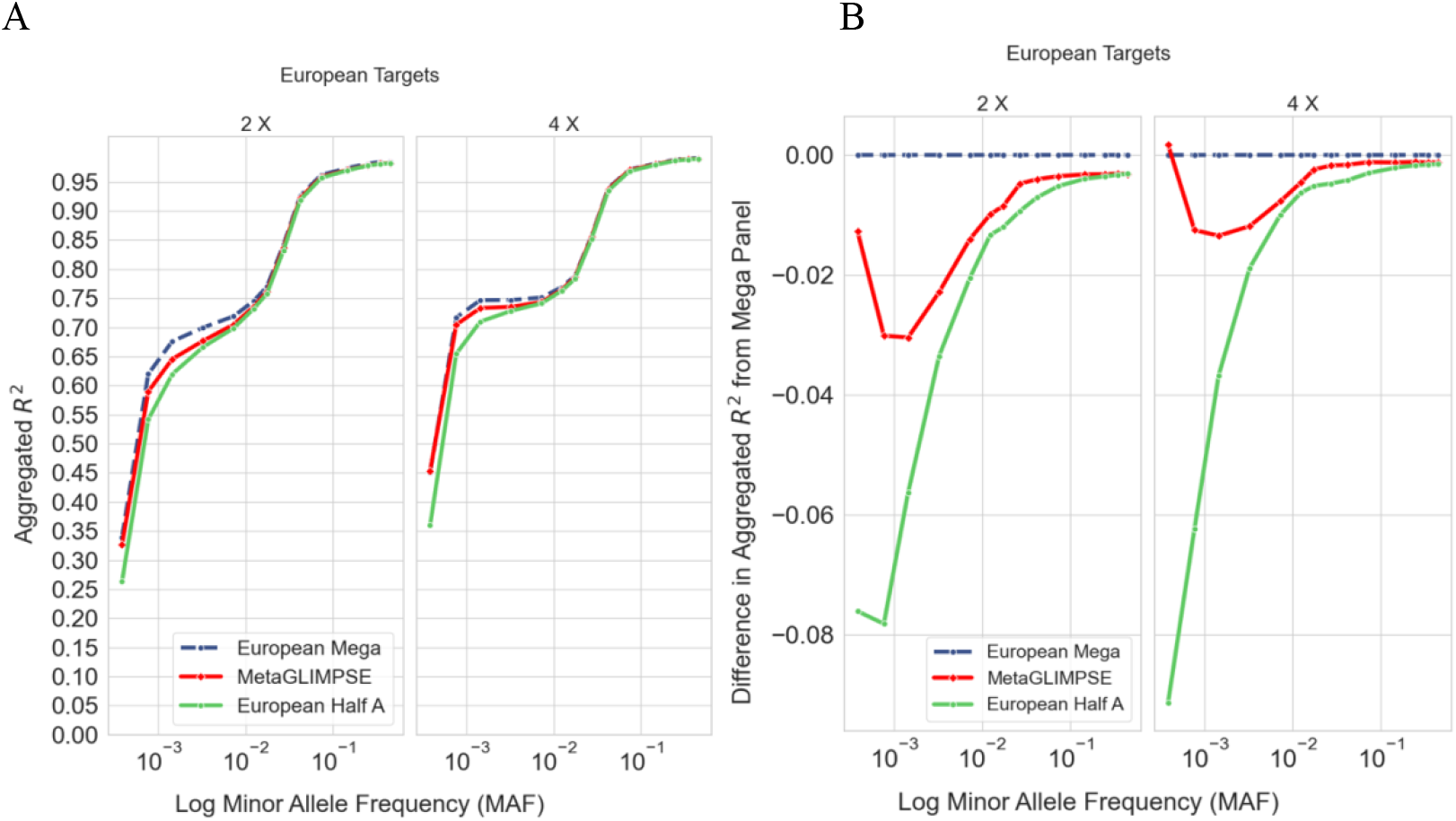
A and B Meta Imputation of European target samples with two European reference panels. We impute 499 Europeans from 1000 Genomes downsampled to 2x and 4x using two European reference panels (n=249). We evaluate MetaGLIMPSE by calculating the aggregated R^2^ between the (meta-)imputed genotypes and true genotypes. We plot both the aggregated R^2^ by MAF (A) and the difference between the aggregated R^2^ from mega imputation and the aggregated R^2^ for each of the other imputations by MAF(B).

At 4x coverage, MetaGLIMPSE’s performance vis a vis mega-imputation improves noticeably compared to 2x. For MAFs between 0.1% and 0.2%, mega-imputation produces an aggregated R^2^ of 0.75 while MetaGLIMPSE delivers 0.73, recovering 62% of the performance gain by mega-imputation compared to the best single-panel imputation (Figure 2a-b).

In general, as coverage increases, MetaGLIMPSE recovers more of the performance gain by mega-imputation, especially for rare variants. At 0.1x, MetaGLIMPSE recovers 25% of the performance increase at MAFs between 0.1% and 0.2% At 8x, for the above MAFs, MetaGLIMPSE recovers 67%. At 8x, for MAF > 1%, the performance of mega-imputation and MetaGLIMPSE are practically equal (Supplementary Figures 3-4).

### Meta Imputing Admixed Samples

Meta-imputation facilitates imputation for admixed populations by combining imputation results from reference panels that each contain a different source population. We model imputation for admixed targets by imputing 57 unrelated African Americans from 1000 Genomes using three imputation panels also from 1000 Genomes: an African population panel without African-Americans (n = 599), a European population panel (n = 499), and a mega-panel (n=1098), which is a concatenation of the African and European reference panels. We generate meta-imputation results by combining the African panel imputation results and European panel imputation results through MetaGLIMPSE and comparing them to mega-imputation and the two single panel imputations.

Across all MAFs and coverages, the mega-imputation has the highest aggregated R^2^ and the European panel nets the lowest aggregated R^2^. Across all coverages, for MAFs > 2%, the performance of MetaGLIMPSE is indistinguishable from the performance of the African panel. For MAFs < 2%, MetaGLIMPSE performs substantially better than the African panel alone. (Supplementary Figures 5-6).

At 2x coverage, for MAFs between 0.5% and 1%, mega-imputation achieves an aggregated R^2^ of 0.62, and MetaGLIMPSE has an aggregated R^2^ of 0.59. The best single panel imputation nets an aggregated R^2^ of 0.58. Across all MAFs and coverages, MetaGLIMPSE is at least as good or better than the best single-panel imputation, which is the African panel. (Figure 3a-b).

**Figure 3.**
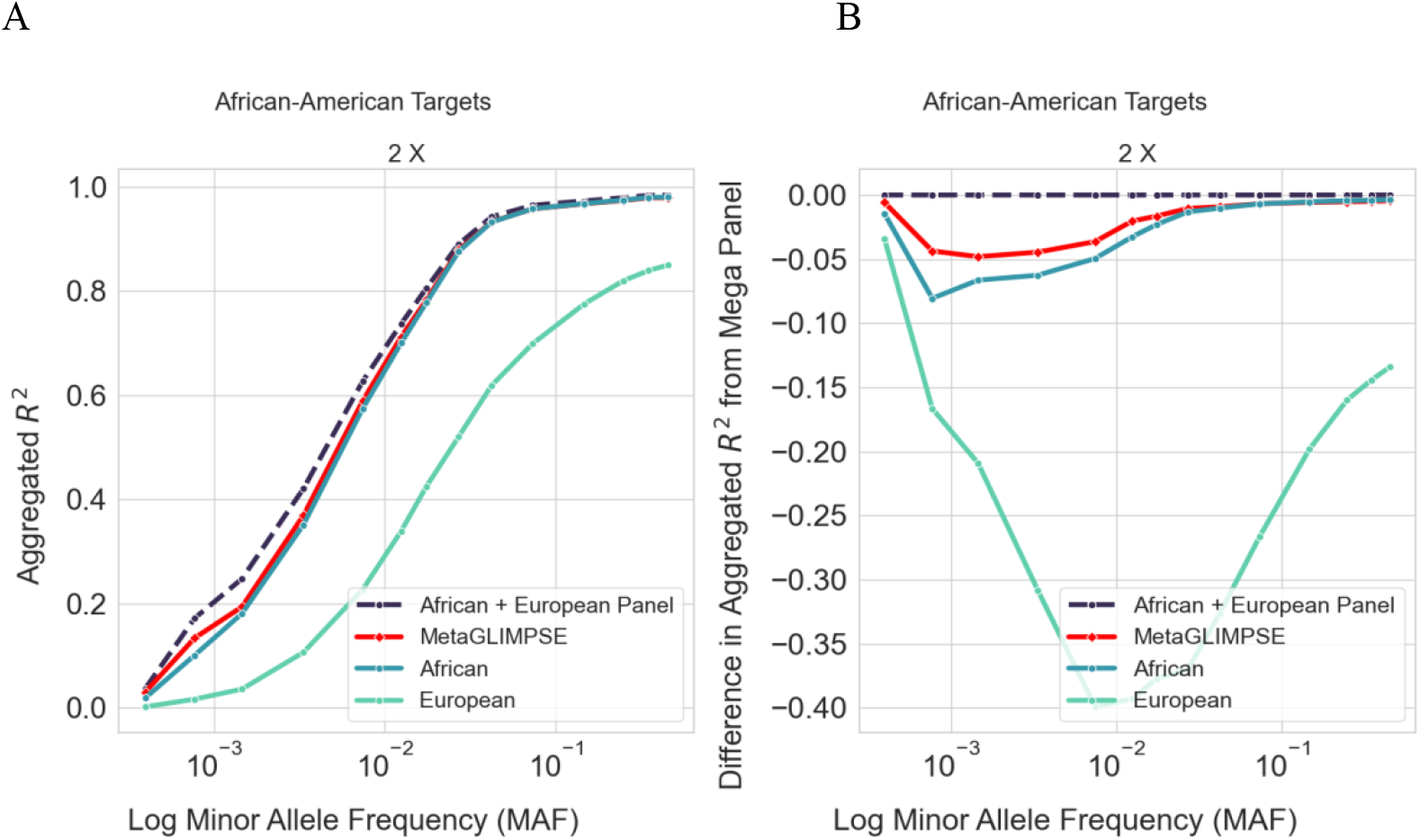
A and B Meta Imputation of African-American target samples with an African and a European reference panel. We impute 57 African-Americans from 1000 Genomes downsampled to 2x using an African (n=599) and a European reference panel (n = 499). We evaluate MetaGLIMPSE by calculating the aggregated R^2^ between the (meta-)imputed genotypes and true genotypes. We plot both the aggregated R^2^ by MAF (A) and the difference between the aggregated R^2^ from mega imputation and the aggregated R^2^ for each of the other imputations by MAF(B).

MetaGLIMPSE’s performance gain compared to the African panel imputation remains constant as coverage increases. For instance, at both 0.1x and 8x, for MAFs between 0.5% and 1%, MetaGLIMPSE recovers 24% of the performance increase by mega-imputation compared to the African panel imputation.

We also find that phasing has an important role in meta-imputation. Across all coverages, when MetaGLIMPSE leverages phasing, it has a higher aggregated R^2^ compared to when it does not– we denote this version MetaGLIMPSE-plain. For example, at 2x, for variants with MAF between 0.5% and 1%, MetaGLIMPSE-plain achieves an aggregated R^2^ of 0.58, which is 0.01 less than MetaGLIMPSE (Supplementary Figures 5-6).

### Meta-Imputing Ancient DNA

MetaGLIMPSE facilitates imputation for aDNA, which is often sequenced at low coverage and imputed with modern day reference panels (Orlando et al 2021; Ausmees et al 2022; Sousa da Mota et al 2023). We emulate this scenario through (meta)-imputing a diverse set of ten target aDNA samples from Sousa da Mota et al. 2023 with the 1000 Genomes African and European panels, as described above. These ten aDNA samples are of European, African, or Native-American ancestry. We evaluate MetaGLIMPSE for these targets by comparing its performance to the mega-panel and the two single-panel imputations.

Across all MAFs and coverages the mega-imputation has the highest aggregated R^2^ followed by MetaGLIMPSE. The single-panel imputations have the lowest aggregated R^2^. The relative performance of each single-panel imputation depends on the MAF (Supplementary Figures 7-8).

At 2x coverage, mega-imputation has the highest aggregated R^2^ across MAFs, and the single-panel imputations have the lowest. Unlike the scenarios with modern targets, mega-imputation has a noticeable performance gain compared to the best single-panel imputation at both rare and common variants. For rare MAFs between 0.2% and 0.5%, MetaGLIMPSE recovers 60% of this performance gain and achieves an aggregated R^2^ = 0.41, while the best single panel imputation drops to 0.27. At common MAFs between 10% and 20%, MetaGLIMPSE has an aggregated R^2^ = 0.92, while the aggregated R^2^ of best single panel imputation drops to 0.91. MetaGLIMPSE recovers 61% of the performance gain by meta-imputation over the best single panel imputation (Figure 5a-b).

**Figure 5.**
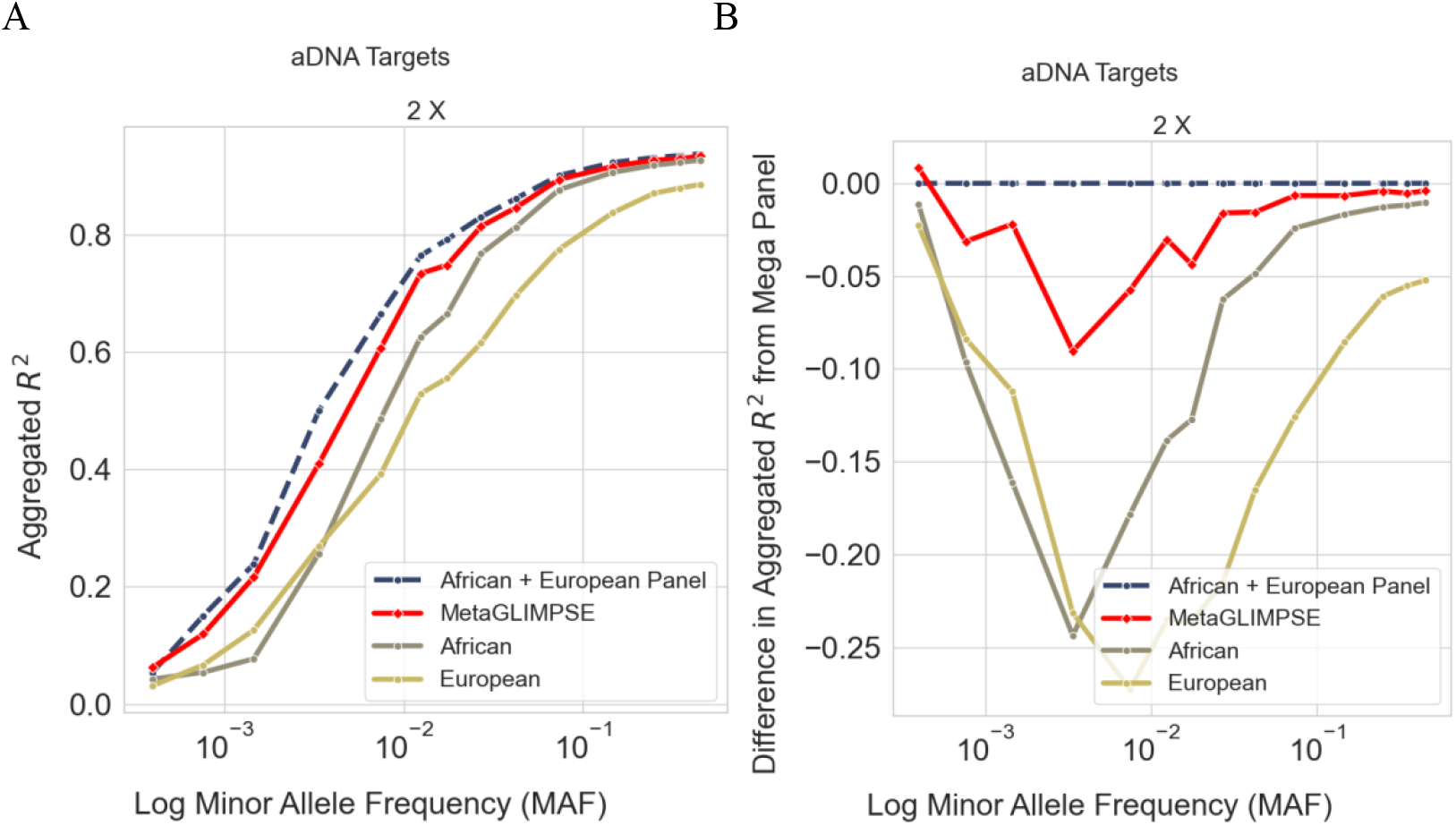
A and B Meta Imputation of aDNA target samples with an African and a European reference panel. We impute 10 aDNA samples used in Sousa da Mota et al. 2023, downsampled to 2x using an African (n=599) and a European reference panel (n = 499). We evaluate MetaGLIMPSE by calculating the aggregated R^2^ between the (meta-)imputed genotypes and true genotypes. We plot both the aggregated R^2^ by MAF (A) and the difference between the aggregated R^2^ from mega imputation and the aggregated R^2^ for each of the other imputations by MAF(B).

As the coverage increases from 0.1x to 8x, MetaGLIMPSE recovers more of the performance gain by mega-imputation over the single panel imputation for rare variants, while staying constant for common variants. At 0.1x, MetaGLIMPSE recovers 36% and 82% of the performance gain at rare variants between 0.2% and 0.5% MAF and common variants between 10% and 20% MAF, respectively. While at 8x, MetaGLIMPSE recovers 71-76% of the performance gain at rare variants with MAF between 0.2% and 0.5% and, at common variants, between 10% and 20% MAF (Supplementary Figures 7-8).

### Computation Time

Computation time is an important consideration in (meta)-imputation algorithms due to the large number of markers processed (Das et al. 2018; Howie et al. 2012; Yu et al. 2022). For HMMs, computation time is linear with the number of markers and quadratic with the number of reference panel pairs. Therefore, for *K* reference panels and *M* markers, the computation time, without file reading and writing, for MetaGLIMPSE is *O(K*^*4*^*M)*, whereas it is *O(K*^*2*^*M)* for MetaGLIMPSE-plain because it has fewer possible configurations for reference panel pairs than MetaGLIMPSE.

We report the computation times for MetaGLIMPSE, including file reading and writing time by the number of samples and markers processed. MetaGLIMPSE computes the underlying HMM in 30,000 marker chunks to allow for parallelization, which significantly speeds up computation time. The number of markers assigned to each chunk is customizable.

MetaGLIMPSE with 1 sample and 900,000 markers takes a total time of 4.42 seconds on an Intel Xeon Platinum 8176 CPU @ 2.10 GHz with 1 core–if we run all 30 chunks in parallel. For 500 samples and ∼900,000 markers, MetaGLIMPSE takes a total time of 32 min and 7 seconds. By comparison, GLIMPSE2 for 500 samples takes 3 hours 15 minutes and 45 seconds. 900,000 markers are roughly the size of the union set of polymorphic markers between the African and European panels in 1000 Genomes on chromosome 20. Since chromosome 20 comprises roughly 2% of the genome, the time for MetaGLIMPSE on a whole genome will be 3 minutes and 41 seconds.

## Discussion

We present a novel meta-imputation framework for low coverage sequencing that enables imputation with multiple reference panels. MetaGLIMPSE is efficient and improves accuracy, achieving consistently better aggregated R^2^ than the best single panel imputation for a wide range of coverages and allele frequencies. This improvement is largest for rare variants, where the performance difference between the mega-imputation and the individual reference panels is also the largest. This is consistent with observations elsewhere that the imputation quality of rare variants is dependent on reference panel size and the extent of shared ancestry with target samples (Das et al. 2018). We observe this improved performance over single-panel imputation for a wide variety of scenarios, including meta-imputing samples with the same ancestry as the reference panels, admixed populations with the source populations as reference panels, and aDNA targets imputed with modern reference panels.

These scenarios model a wide range of applications. First, the scenario of meta-imputing targets with the same ancestry as the reference translates to combining imputation results for e.g. European target samples from several reference panels such as TopMed and HRC that cannot be used in a mega-imputation framework for logistic reasons. The results are relevant for e.g. meta-imputing European targets sequenced using blended genome exome (BGE) (Boltz et al. 2024). Since BGE is sequenced at 1-4x coverage (Boltz et al. 2024), our results suggest that meta-imputation of the low-coverage regions would be nearly equal to mega-imputation. Second, our scenario of imputing admixed samples with the source populations separately indicates that MetaGLIMPSE would facilitate imputing admixed samples, such as those in the Mexican Biobank. There, the best reference panel for imputing a given sample depends on its global ancestry composition (Ziyatdinov et al. 2023). Our results illustrate that combining imputation results will never reduce the quality of the imputation and so meta-imputation with MetaGLIMPSE is a powerful strategy if the best reference panel is unknown or if different panels are better in different parts of the genome. Third, the scenario of imputing ancient DNA from samples that are spatially and temporally diverse using two modern reference panels from different ancestries, directly applies to aDNA imputation, for which the available reference panels consist of modern humans (Allentoft et al. 2024; Ringbauer et al. 2024; Sousa da Mota et al 2023). Meta imputing aDNA with 1000 Genomes as well as a population specific panel more closely related to the target sample could improve the imputation accuracy of aDNA. For instance, for ancient European targets, this additional reference panel could be UK Biobank 200K (Ribeiro et al. 2023), which is publicly available to anyone approved to use UK Biobank data.

Our results also encourage creating an aDNA reference panel. As of now, aDNA is imputed with modern reference panels, most commonly, 1000 Genomes (Allentoft et al. 2024; Ringbauer et al. 2024; Sousa da Mota et al. 2023). The current literature on imputing aDNA (Allentoft et al. 2024; Ausmees et al. 2022; Ausmees et al. 2023; Biddanda et al. 2022; Hui et al. 2020; Sousa da Mota et al. 2023) does not exploit the 10,000+ and growing set of aDNA samples (Mallick et al. 2024), from which one could create an aDNA reference panel. Meta imputing aDNA with both a small aDNA-only panel and a large modern panel would leverage these aDNA samples. Due to their lower coverage and higher genotyping error rate (Ausmees et al. 2023; Hui et al. 2020; Garrido Marques et al. 2024) than modern reference panel genomes, aDNA might not be selected as the best template haplotype by imputation algorithms. MetaGLIMPSE, which explicitly allows for aDNA as templates in imputation algorithms, could provide a substantial benefit– especially for older ancient genomes, which have undergone many subsequent population replacements (Orlando et al. 2021).

Our method is easily adaptable to other low coverage sequencing imputation algorithms other than GLIMPSE2, such as Beagle 4.1 (Browning et al. 2016) and QUILT (Davies et al. 2021). Furthermore, MetaGLIMPSE could also meta-impute samples imputed with multiple low coverage sequencing algorithms. Some algorithms impute certain variants better than others (Rui et al. 2021), necessitating meta-imputation.

We made several choices during our analysis. First, we do not present meta-imputation with large reference panels, such as HRC and TopMed, since they are not publicly available for imputation with low coverage sequencing data. Therefore, many low coverage sequencing studies use 1000 Genomes as their reference panel (Allentoft et al. 2024; Santos et al. 2024; Sousa da Mota 2023). While we do not explicitly model the size of the reference panel in MetaGLIMPSE, we expect that since our method takes a weighted average of single panel imputation estimates, it should extend to larger reference panels. Second, we handle variants missing in only one of the two reference panels by setting their genotype to the reference. Deciding how to handle variants that are not polymorphic in all reference panels is a challenging consideration for meta-imputation (Yu et al. 2022). Setting the missing variant to the reference is correct in the vast majority of cases. However, individuals of the same ancestry as those in the reference panel that have a non-reference allele might not have been sequenced. Future work would involve constructing a more robust strategy to handle such variants.

In summary, we propose a new method to combine low-pass imputation results that consistently outperforms the best single panel imputation for a wide range of scenarios—including target samples and reference panels from the same ancestry, admixed target samples with reference panels from each source population, and aDNA target samples with modern reference panels. MetaGLIMPSE is especially useful for increasing imputation accuracy of rare variants, and we expect our method will help increase the imputation accuracy for samples of diverse ancestries and time periods by allowing for privacy protected imputation with multiple reference panels.

## Code and Data Availability

The code for MetaGLIMPSE along with tutorial and instructions is publicly available at https://github.com/karinkumar/MetaGLIMPSE. The links to the software used are as follows: bwa-mem2 (https://github.com/bwa-mem2/bwa-mem2), bcftools and samtools (https://www.htslib.org/), and GLIMPSE2 branch ap-option (https://github.com/odelaneau/GLIMPSE/tree/ap-field). This branch of GLIMPSE2 provides the inputs necessary to run MetaGLIMPSE. The 2504 samples from Phase 3 1000 Genomes resequenced at 30x are available at the European Nucleotide Archive: PRJEB31736. The aDNA samples originate from the following studies: SIII (https://doi.org/10.1126/science.aao1807), baa001, ela001, new001 (https://doi.org/10.1126/science.aao6266), Lovelock2 and Lovelock3 (https://doi.org/10.1126/science.aav2621), and atp016 (https://doi.org/10.1073/pnas.1717762115) and STT-A-2, SSG-A-2, HSJ-A-1 (https://doi.org/10.1126/science.aar2625).

## Acknowledgements

This work was funded by the NIH Grants R01 HG011031 and R01 HG005855 to S.Z.. We acknowledge Ketian Yu and Jean Morrison for helpful conversations and support. We also thank the individuals whose genomic data we used in our analyses.

## Author Contributions

K.H.K. and S.Z. developed the ideas. K.H.K. and S.Z. wrote the manuscript with contributions from S.R., who also developed the inputs for the software. K.H.K. implemented the software and performed the analyses.

