## Supplement for "MetaGLIMPSE: Meta Imputation of Low Coverage Sequencing Data for Modern and Ancient Genomes"

### Supplemental Text

*Meta Imputing Populations Absent from Reference Panels*

We consider a scenario that is applicable to a sequenced population that either does not exist in any reference panel or is not in an accessible reference panel. We mimic this scenario by imputing a target population using two reference panels that are not closely related to the target population. Specifically, we meta-impute 504 East Asians from 1000 Genomes with the same African and European reference panels described in the main text. We compare MetaGLIMPSE to the mega-imputation and the two single panel imputations.

For coverages between 0.5x and 8x, mega imputation has the highest aggregated $R^2$ followed by MetaGLIMPSE, the European panel imputation and, finally, the African panel imputation, respectively. Across all coverages, the aggregated $R^2$ is overall much smaller for lower frequency alleles (MAF < 1%) than in the other scenarios. (Supplementary Figures 9-10).

At coverages of 0.5x and greater, MetaGLIMPSE practically matches the aggregated $R^2$ of the mega-imputation across all MAFs. For instance, at 2x, for MAFs between 3.5% and 5%, MetaGLIMPSE matches the aggregated $R^2$ of the mega-imputation (aggregated $R^2$ = 0.76) (Supplementary Figure 1a-b). This result is consistent across coverages. At 0.5x and 8x, for the above MAFs, MetaGLIMPSE has the same aggregated $R^2$ as the mega-imputation at values of 0.60 and 0.86, respectively (Supplementary Figures 9 -10).

A

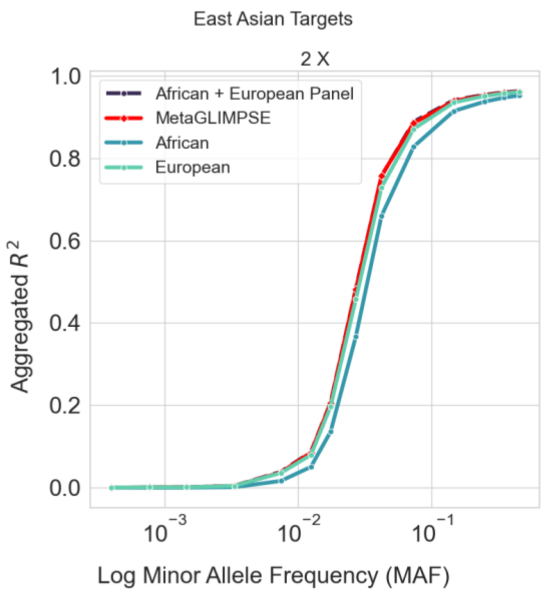


B

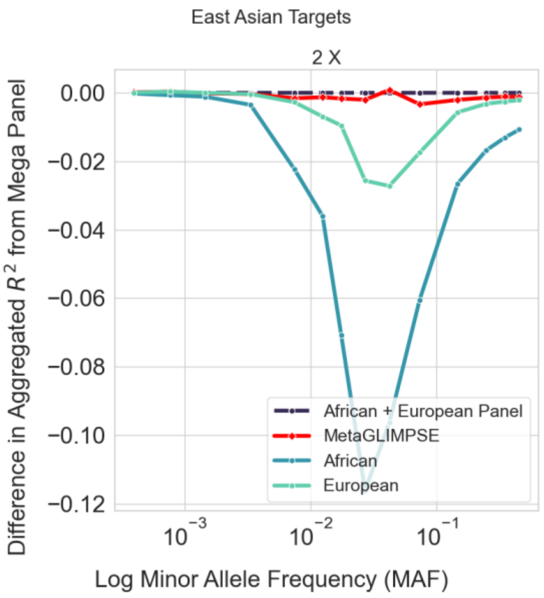

**Figure S1A and B Meta Imputation of East Asian target samples with an African and a European reference panel.** We (meta-)impute 504 East Asians from 1000 Genomes downsampled to 2x using an African (n=599) and a European reference panel (n = 499). We evaluate MetaGLIMPSE by calculating the aggregated $R^2$ between the (meta-)imputed genotypes and true genotypes. We plot both the aggregated $R^2$ by MAF (A) and the difference between the aggregated $R^2$ from mega imputation and the aggregated $R^2$ for each of the other imputations by MAF(B).

*Meta-Imputing at Ultra Low Coverages*

We consider meta-imputation at ultra-low coverages (e.g. 0.1x), which are common in fields like aDNA (Orlando et al. 2021). For the scenarios where (i) both reference panels are from the same ancestry as the target, (ii) where the reference panels are the source populations to an admixed target, and (iii) where the targets are aDNA and the reference panels are modern, the patterns for 0.1X are consistent with the patterns for higher coverage: the mega-imputation is most accurate, followed by meta-imputation and then the individual panels (Supplementary Figures 3 - 8). However, in the scenario where both reference panels are selected from different populations than the target, the results at 0.1x differ from the patterns observed at higher coverages (Supplementary Figures 1a-b; 2a-b).

For this scenario, the relative performance of the different imputations varies by MAF. For MAFs less than 1.5%, MetaGLIMPSE achieves an aggregated $R^2$ comparable to the best single-panel imputation– the European panel imputation– which the mega-imputation performs equal to or worse than. For MAFs between 1.5% and 5%, MetaGLIMPSE achieves an equal or higher aggregated $R^2$ than the European panel imputation and the mega-imputation, which performs worse than the European panel imputation. For MAFs higher than 5%, MetaGLIMPSE outperforms the mega-panel imputation, but not the European panel imputation.

A B

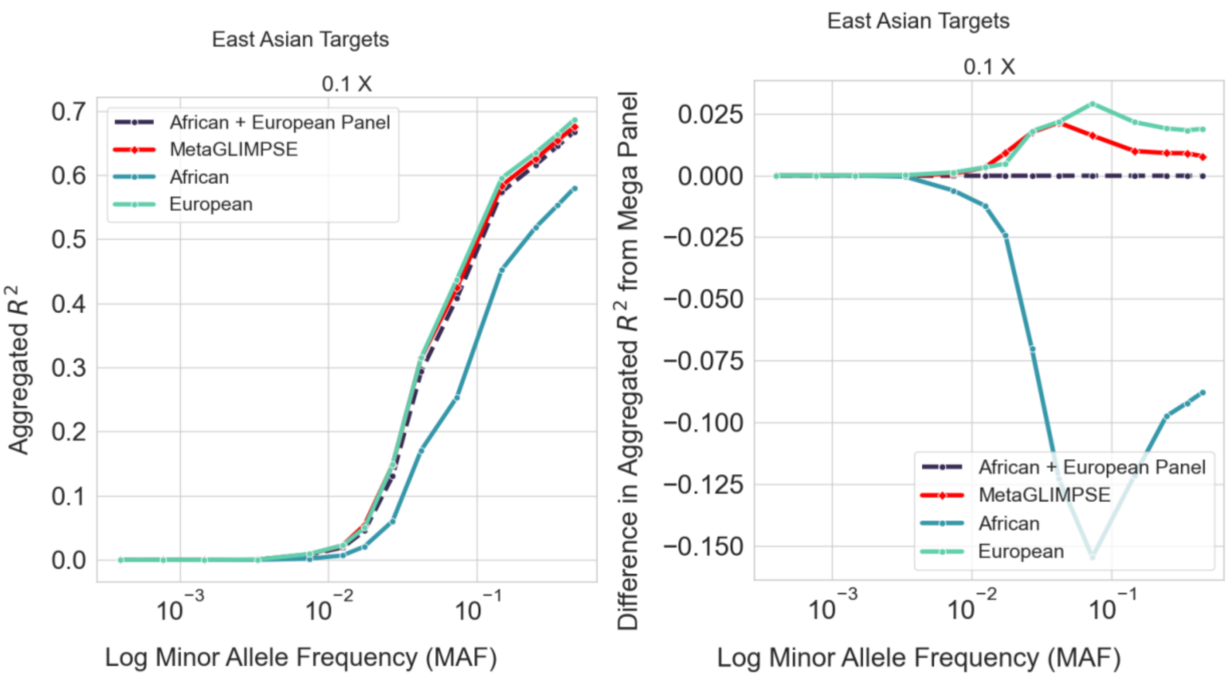


**Figure S2A and S2B Meta Imputation of East Asian target samples at 0.1x with an African and a European reference panel.** We (meta-)impute 504 East Asians from 1000 Genomes downsampled to 0.1x using an African (n=599) and a European reference panel (n = 499). We evaluate MetaGLIMPSE by calculating the aggregated $R^2$ between the (meta-)imputed genotypes and true genotypes. We plot both the aggregated $R^2$ by MAF (A) and the difference between the aggregated $R^2$ from mega imputation and the aggregated $R^2$ for each of the other imputations by MAF(B).

### Supplemental Tables

| Sample | Reported Genome-wide Coverage | Coverage on hg38 Chr 20 | C→T | Ancestry | Age (years before present) | Publication |
|---|---|---|---|---|---|---|
| atp016 | 13.2 | 13.9 | 0.185 | European | 4867-5212 | Valdiosera et al., PNAS (2018) |
| baa001 | 13.5 | 17.1 | 0.045 | African | 1831-1986 | Schlebusch et al., Science (2017) |
| ela001 | 13.5 | 13.5 | 0.030 | African | 453-533 | Schlebusch et al., Science (2017) |

| | | | | | | |
|---|---|---|---|---|---|---|
| new001 | 10.9 | 14.6 | 0.037 | African | 327-508 | Schlebusch et al., Science (2017) |
| SIII | 11.2 | 11.3 | 0.016 | European | 33031-35154 | Sikora et al., Science (2017) |
| Lovelock2 | 15.4 | 15.4 | 0.008 | Native American | 1818-1942 | Moreno-Mayar et al., Science (2018) |
| Lovelock3 | 19.1 | 19.2 | 0.008 | Native American | 567-687 | Moreno-Mayar et al., Science (2018) |
| SSG-A-2* | 10.2 | 16.7 | NA | European | 950-1100 | Ebenesersdottir et al., Science (2018) |
| STT-A-2* | 14.1 | 25.7 | NA | European | 950-1050 | Ebenesersdottir et al., Science (2018) |
| HSJ-A-1* | 29.4 | 41.8 | NA | European | 950-1080 | Ebenesersdottir et al., Science (2018) |

*bams already aligned to hg38

Supplementary Table 1: Summary statistics of aDNA targets used to evaluate MetaGLIMPSE

**Supplementary Figures**

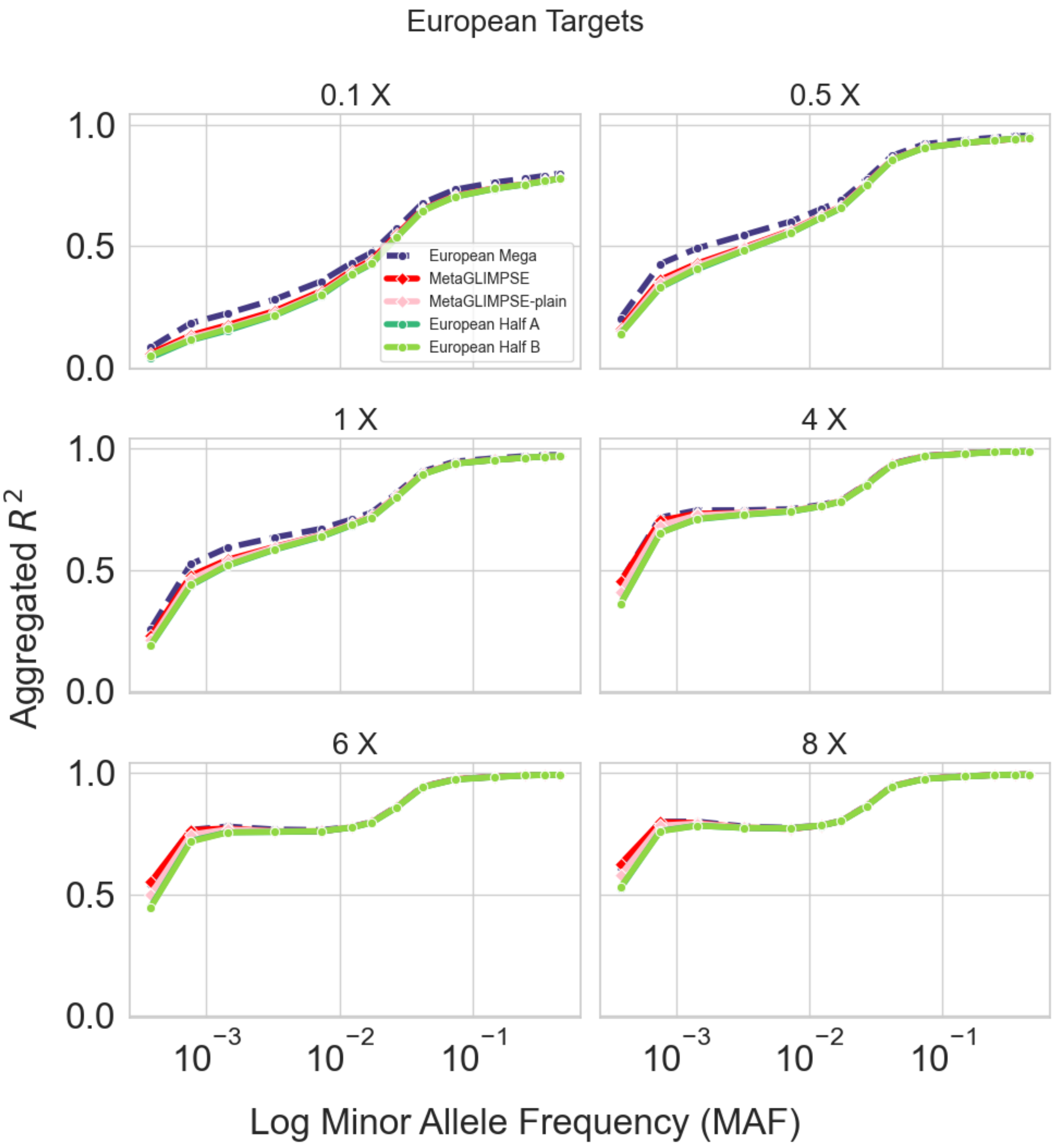

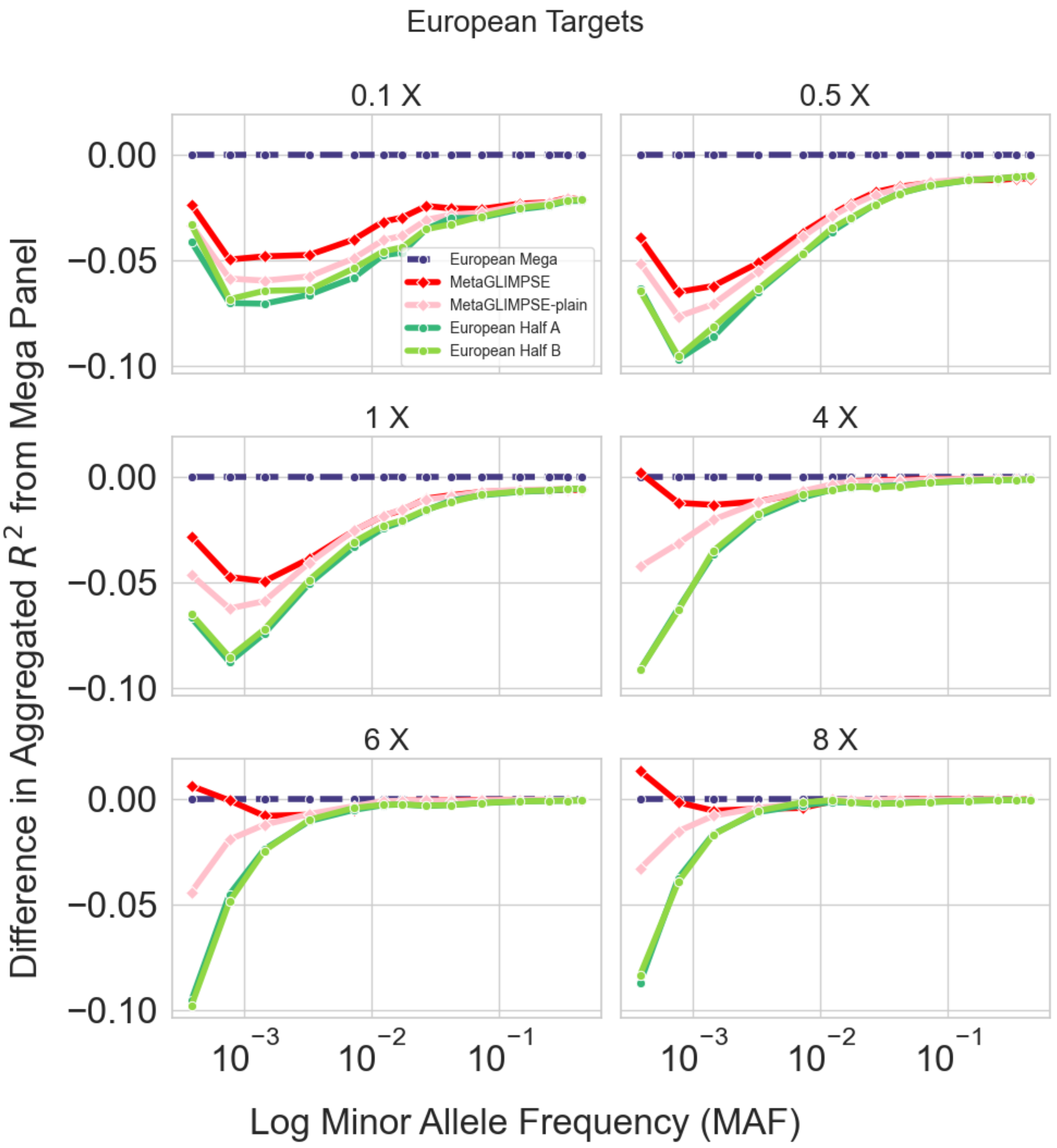


**Figures S3 and S4: Meta-Imputation with European Panels for European Target Samples Across Coverages 0.1x - 8x.** We (meta-)impute 499 Europeans from 1000G downsampled from 0.1x to 8x using two European reference panels (n=249). We evaluate MetaGLIMPSE by meta-imputing the results and calculating the aggregated $R^2$ between the (meta-)imputed genotypes and true genotypes. We plot the aggregated $R^2$ for each imputation by MAF. We report the aggregated $R^2$ (A) and the difference between the aggregated $R^2$ from mega imputation and the aggregated $R^2$ for each of the other imputations (B).

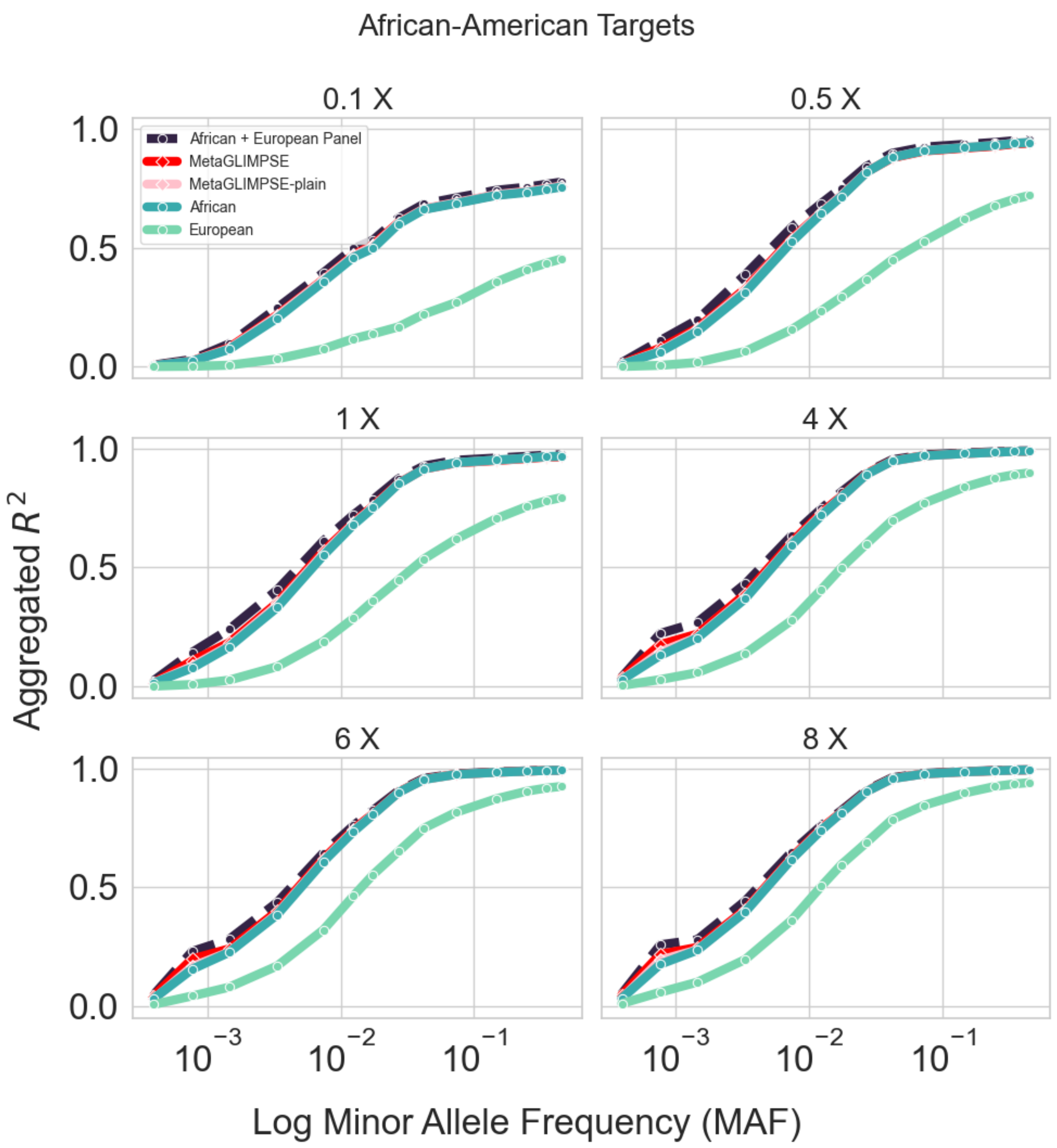

African-American Targets
0.1 X
0.5 X
1 X
4 X
6 X
8 X
African + European Panel
MetaGLIMPSE
MetaGLIMPSE-plain
African
European
1.0
0.5
0.0
Aggregated $R^2$
$10^{-3}$
$10^{-2}$
$10^{-1}$
Log Minor Allele Frequency (MAF)


132

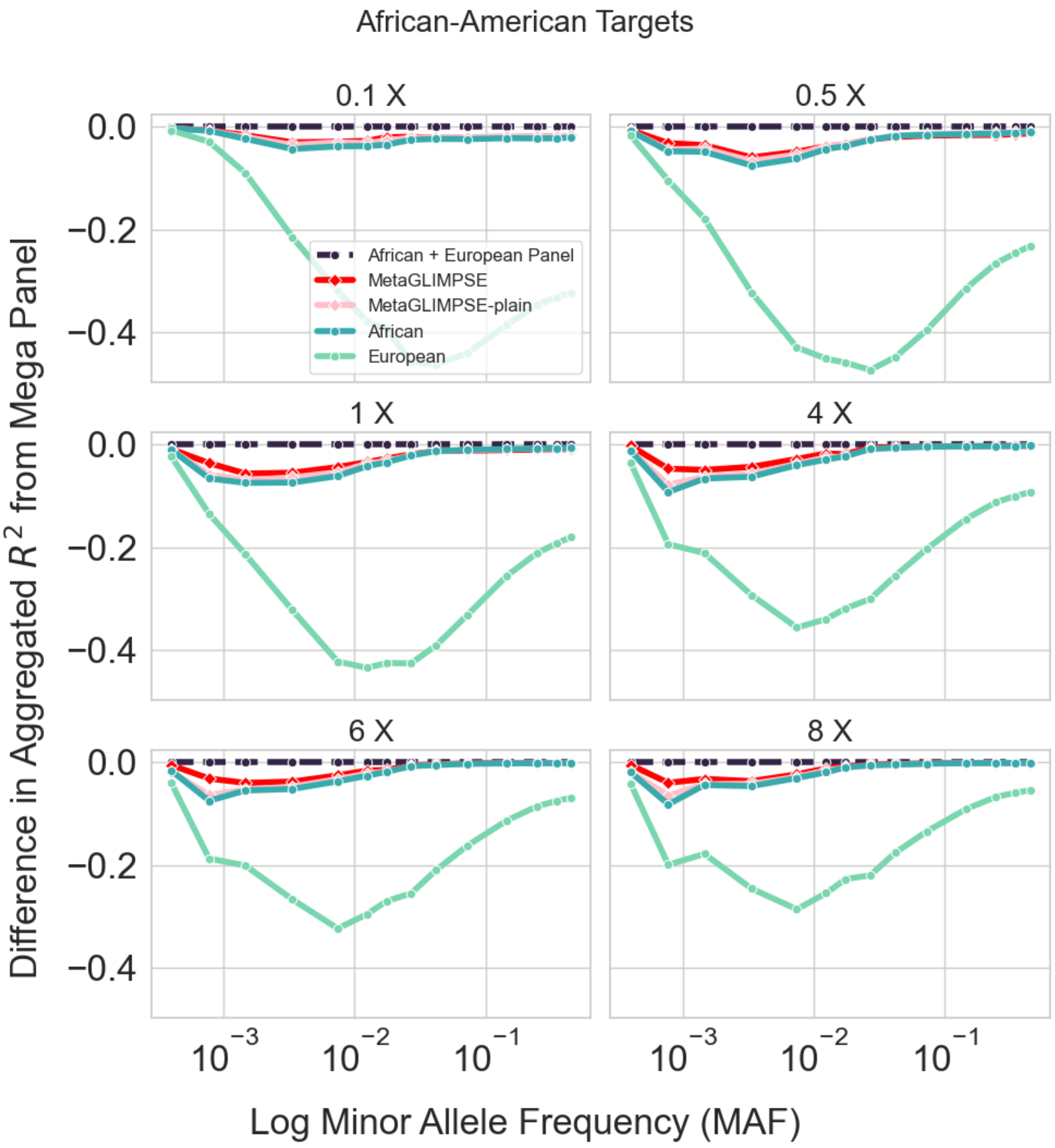


**Figures S5 and S6 Meta-Imputation for African-American Target Samples with African, European Panels Across Coverages 0.1x - 8x.** We (meta-)impute 57 African-Americans from 1000G downsampled from 0.1x to 8x using an African (n=599) and a European reference panel (n = 499). We evaluate MetaGLIMPSE by meta-imputing the results and calculating the aggregated $R^2$ between the (meta-)imputed genotypes and true genotypes. We plot the aggregated $R^2$ for each imputation by MAF. We report the aggregated $R^2$ (A) and the difference between the aggregated $R^2$ from mega imputation and the aggregated $R^2$ for each of the other imputations (B).

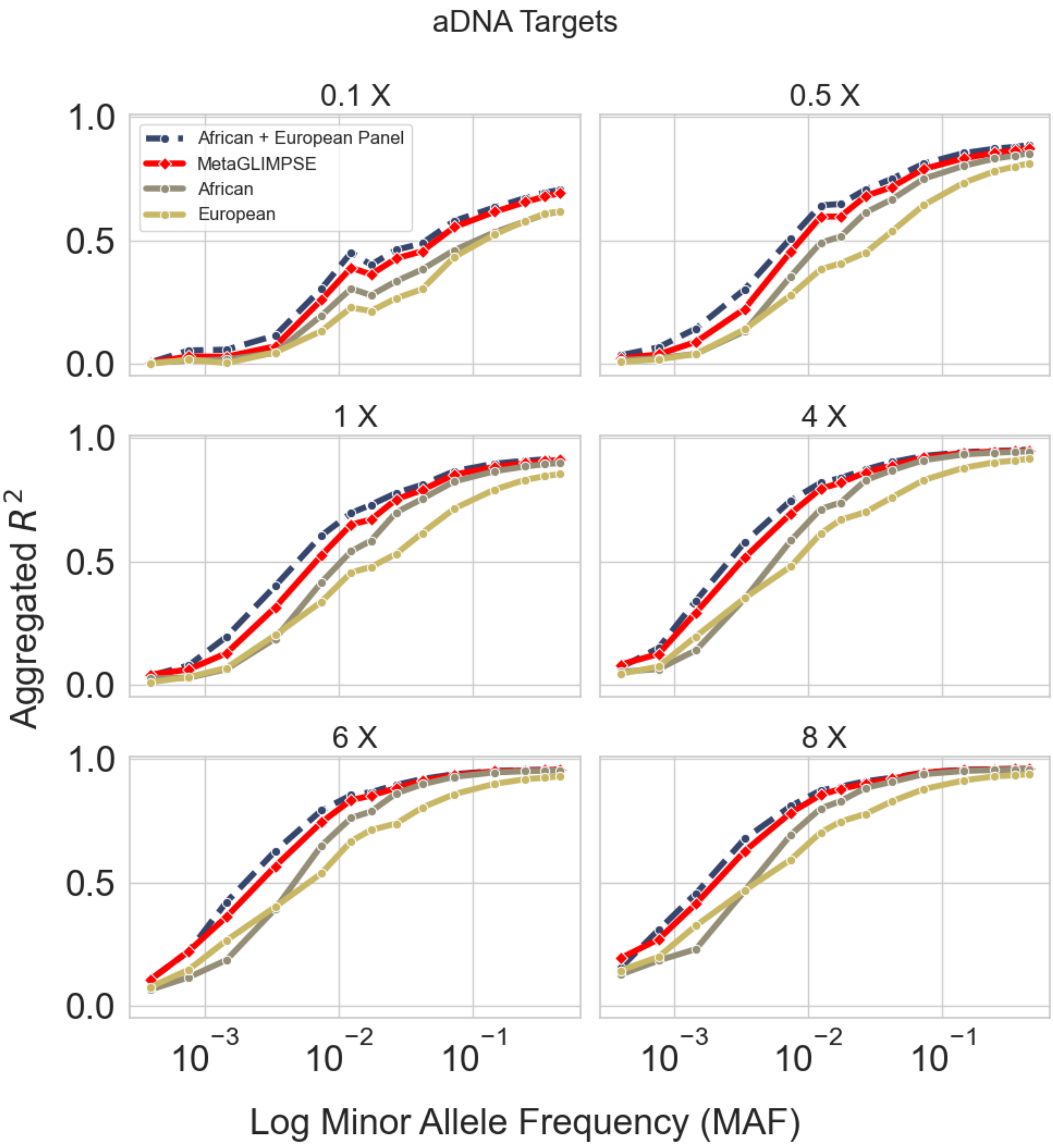
aDNA Targets
0.1 X
0.5 X
1 X
4 X
6 X
8 X
African + European Panel
MetaGLIMPSE
African
European
Aggregated $R^2$
Log Minor Allele Frequency (MAF)
1.0
0.5
0.0
$10^{-3}$
$10^{-2}$
$10^{-1}$

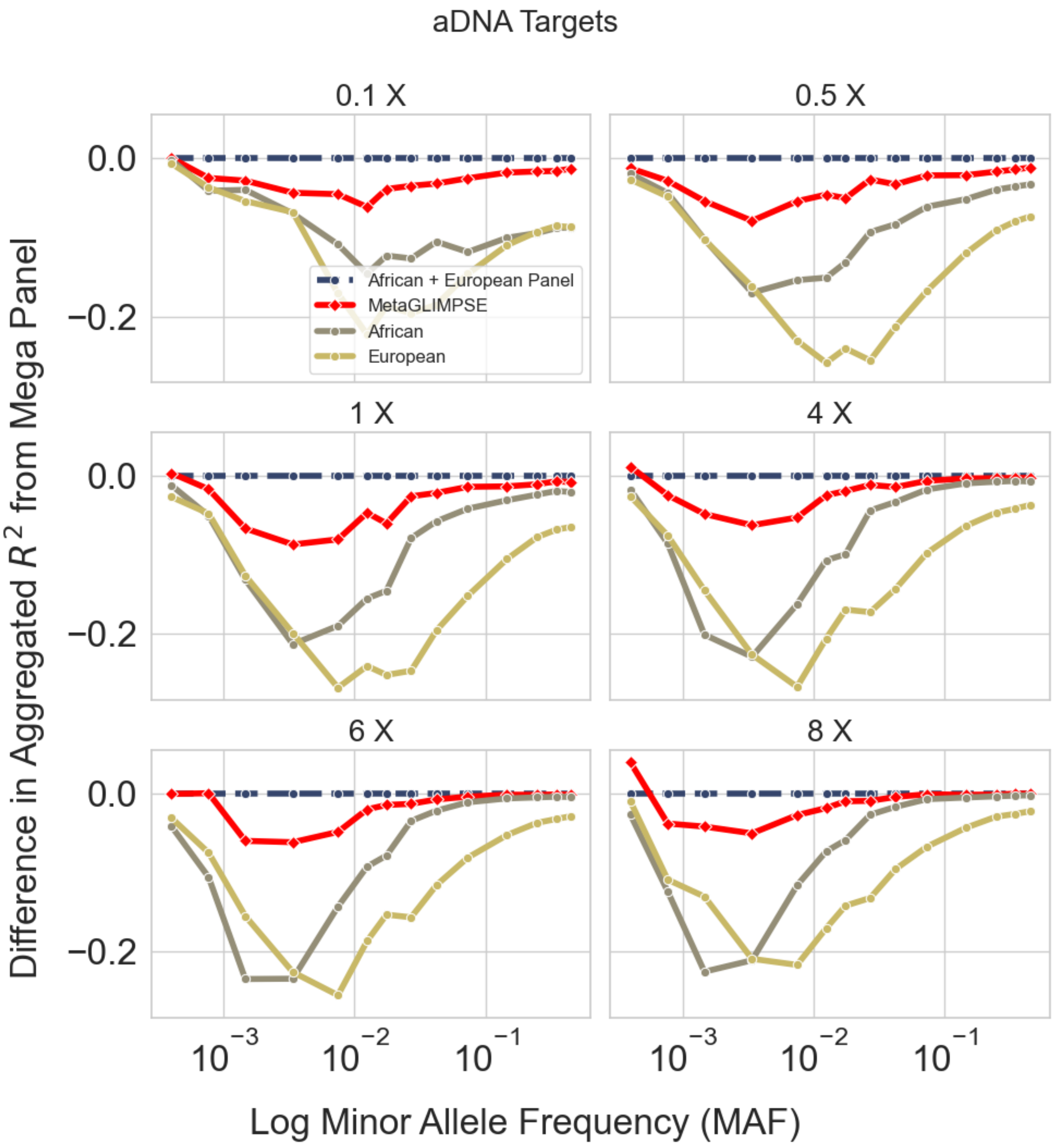


**Figures S7 and S8 Meta-Imputation for aDNA Target Samples with African, European Panels Across Coverages 0.1x - 8x.** We (meta-)impute 10 aDNA samples from Sousa da Mota et al. 2023 downsampled from 0.1x to 8x using an African (n=599) and a European reference panel (n = 499). We evaluate MetaGLIMPSE by meta-imputing the results and calculating the aggregated $R^2$ between the (meta-)imputed genotypes and true genotypes. We plot the aggregated $R^2$ for each imputation by MAF. We report the aggregated $R^2$ (A) and the difference between the aggregated $R^2$ from mega imputation and the aggregated $R^2$ for each of the other imputations (B).

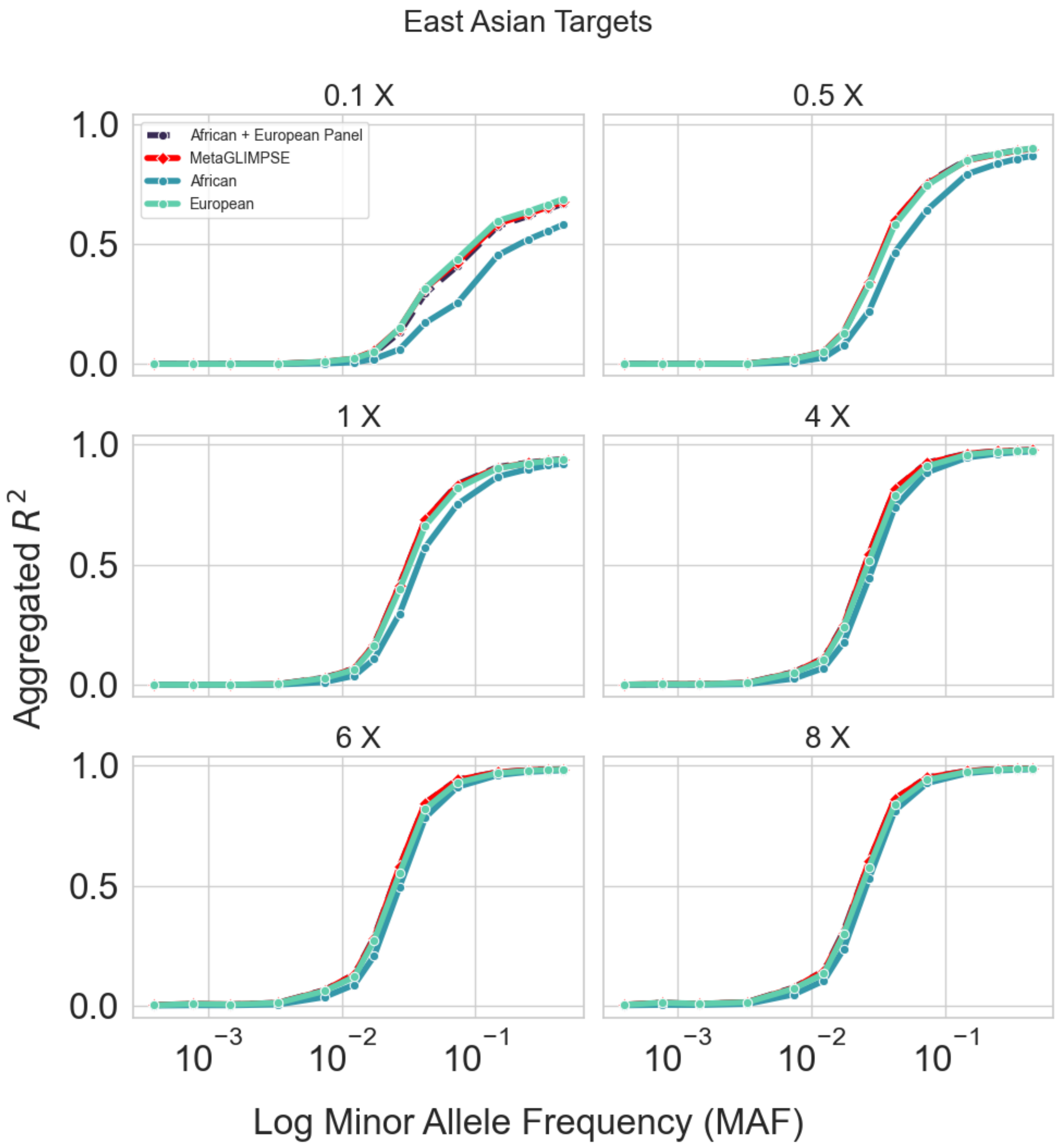
East Asian Targets
0.1 X
0.5 X
1 X
4 X
6 X
8 X
African + European Panel
MetaGLIMPSE
African
European
Aggregated $R^2$
1.0
0.5
0.0
$10^{-3}$
$10^{-2}$
$10^{-1}$
Log Minor Allele Frequency (MAF)

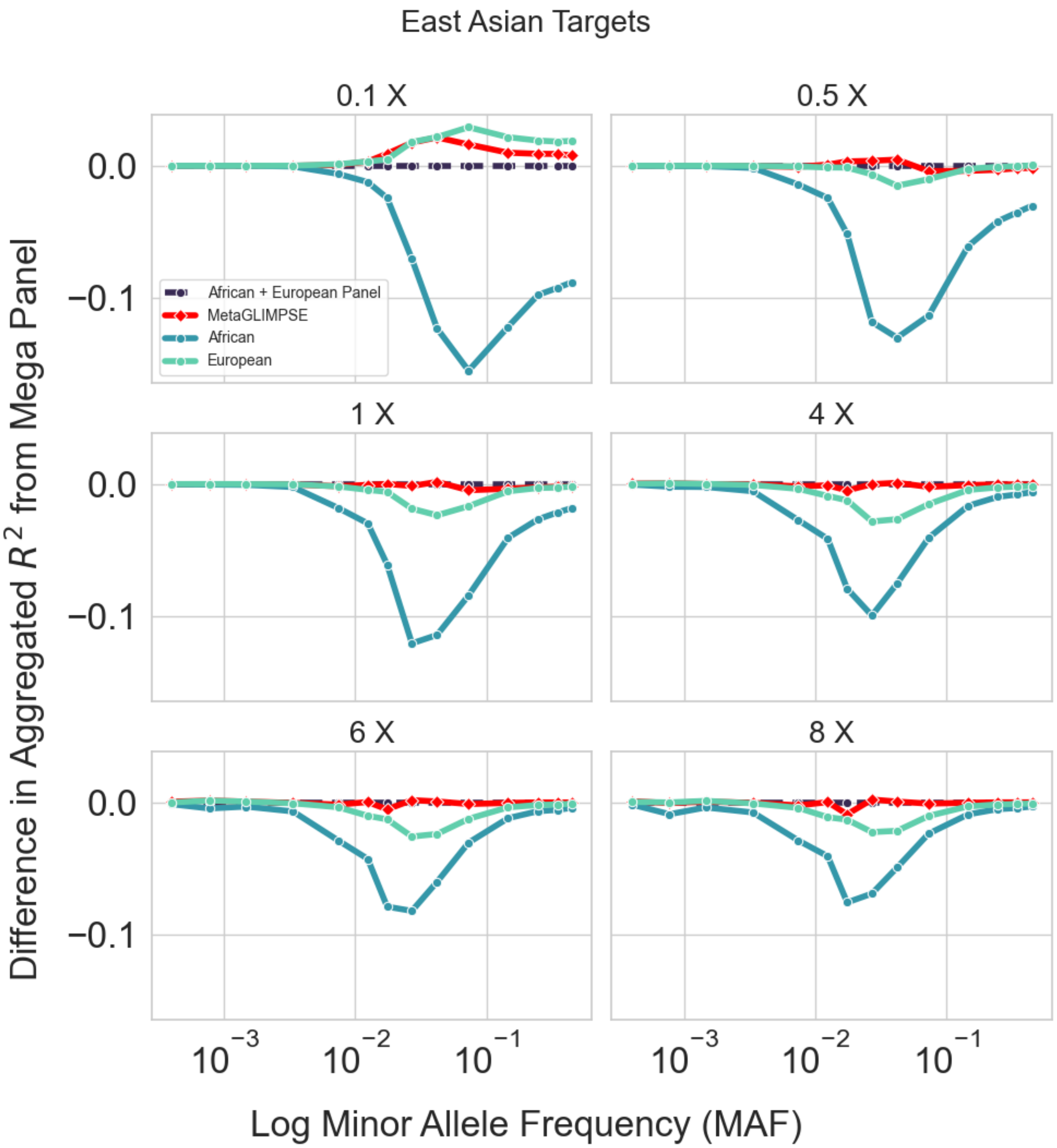


**Figures S9 and S10 Meta-Imputation for East Asian Target Samples with African, European Panels Across Coverages 0.1x - 8x.** We (meta-)impute 504 East Asians from 1000G downsampled from 0.5x to 8x using an African (n=599) and a European reference panel (n = 499). We evaluate MetaGLIMPSE by meta-imputing the results and calculating the aggregated $R^2$ between the (meta-)imputed genotypes and true genotypes. We plot the aggregated $R^2$ for each imputation by MAF. We report the aggregated $R^2$ (A) and the difference between the aggregated $R^2$ from mega imputation and the aggregated $R^2$ for each of the other imputations (B).

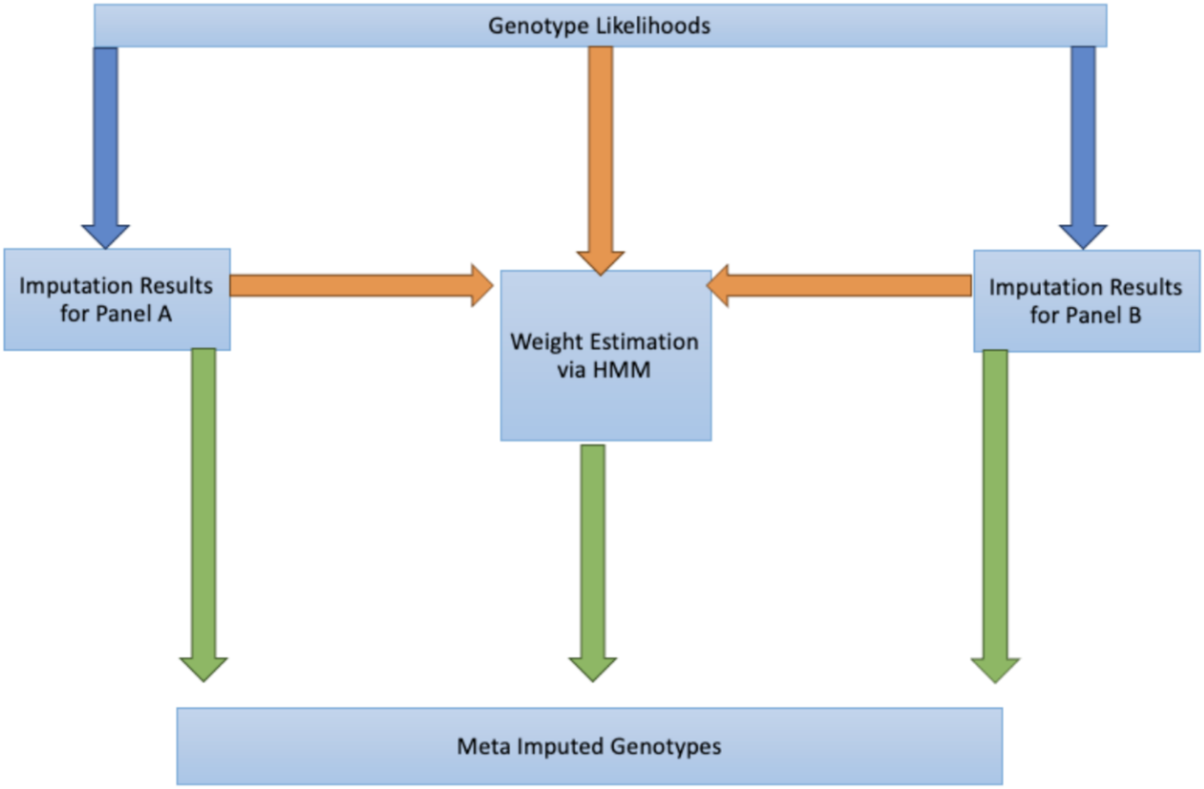


**Figure S11 MetaGLIMPSE workflow.** The workflow for MetaGLIMPSE starts with imputing the target samples with the single reference panels using the genotype likelihoods calculated from the sequence data, denoted by the blue arrows. Here we show two panels: A and B; however, MetaGLIMPSE can combine any number of reference panels. The second step is to estimate weights customized for each individual and marker using a HMM, denoted by the orange arrows. The third and final step is to generate meta imputed genotypes by weighting each of the imputation estimates, denoted by the green arrows.
